# Preliminary Survey of Fungal Communities Across a Plastics/No Plastics Transition on an Oregon Beach

**DOI:** 10.1101/2024.01.18.576272

**Authors:** Ken Cullings, Karisa Boyce Arterbury, Richard Arterbury

**Affiliations:** Ocean Blue Project, 6699 Fox Centre Pkwy Unit 104-146, Gloucester, VA 23061

## Abstract

Plastics pose an increasing and significant threat to both human and environmental health. While many fungi can degrade a variety of organic polymers, investigations into which fungi possess the potential to remediate environmental plastics contamination have only recently become a priority. To help address this need, we tested the null hypothesis that chronic plastics contamination has no impact on the fungal communities across a plastics/no plastics transition in a beach sand in northern Oregon. We used sieving and binocular microscopy of microplastics (particle size, 12.6µm +/-5.5µm, detection range 1-5000µm) to confirm the plastics/no plastics transition. We used paired plot design to collect samples across this transition and analyzed the fungal communities using high-throughput DNA sequencing methods for fungal ITS-2. Results indicated that the beach sand contaminated with plastics held an extensive fungal community, while un-contaminated sand held no fungal community at all. System dominants included *Acremonium* and *Penicillium*, both free-living ascomycete fungi that have shown plastics-degrading capabilities in lab studies, and the ectomycorrhizal genus, *Russula* a symbiotic fungus that has known plastics-degrading enzyme capabilities. Also amongst dominant genera was a human fungal pathogen (genus *Malassezia*) that causes chronic skin disease. These results provide new fungal models for further studies of fungal and ectomycorrhizal remediation of plastics contaminated contaminated beach sand.

## Introduction

Plastics and microplastics represent a persistent and growing global environmental health problem (Reviewed by 1) Hence, managing this problem is of paramount importance. To date, landfilling remains the most often used method of disposal and though methods such as incineration and recycling are both expensive and environmentally hazardous, and hence lack the scale to provide a sufficient solution. In light of this, focus has been shifting to the use of microbes, including fungi, as vehicles for plastics degradation and elimination.

Fungi are widespread and resilient, inhabiting both terrestrial and aquatic habitats, and function as the Earth’s recyclers able to break down some of the most recalcitrant polymers, including plastics (1). Research is showing that some fungi may possess the capability in the lab, not all fungi are adapted to conditions necessary to perform the functions in nature or under conditions created by industrial processes (e.g. 2). Thus, using metagenomic methods to identify fungi in the “plastisphere” that could be effective either as free-living degraders in plastics-polluted areas and/or as sources of enzymes for use in engineered recycling methods is of paramount importance (1). In this study we performed a preliminary survey of the fungal communities across a plastics/no plastics transition in a beach sand habitat in Oregon in order to ascertain impacts of plastics contamination on fungal communities in this habitat. The overall goal of our work is to not only study impacts of plastics contamination on beach sands, but also to locate fungal candidates for use in plastics degradation and remediation.

## Methods

### Collections

Samples were taken at a beach in Oregon (45.722568, -123.941561) across a transition of sand containing no plastics to sand chronically contaminated by plastics. To confirm the state of contamination in the two habitat types, three samples from each of three plots in each sand type were collected into sterile 50ML Falcon tubes and sent to EMSL Analytical, Inc in Cinnaminsom NJ for analysis by sieve separation and stereoscopic microscopic analysis, particle size range 1µm -5000µm diameter.

### Molecular Methods

Three samples from each of three plots on each side of the transition were collected into sterile 50ML Falcon tubes and sent to Novogene Corp for analysis. The fungal microbiomes of the three beach sand types were determined via Internal Transcribed Spacer (ITS-2) amplicon sequencing (itags) using fungal-specific primers (ITS3F; GCATCGATGAAGAACGCAGC-ITS4R; TCCTCCGCTTATTGATATGC). PCR reactions were undertaken using sand extractions using the PHusion High-Fidelity PCR Master Mix (New England Biolabs). Libraries were generated using the TruSeq DNA PCR-Free Prep Kit (Illumina) and quality was assessed using Qubit 2.0 on a Thermo Scientific Fluorometer and Agilent Bioanalyzer 2100 system. The library was sequenced using an IlluminHiSeq2500 platform. ITS-2 itags were generated by Novogene Corporation. Sequences were assembled using FLASH V1.2.7 (3) and data were quality filtered using QIIME V2 using the default parameters (4). Chimeras were removed using UCHIME (5). Sequences were analyzed using UPARSE v7.0.1001 (6) and sequences with >97% similarity were clustered as OTUs. Multiple sequence alignments were performed using MUSCLE V3.8.31 (7). Taxonomic annotation was accomplished using the GreenGene Database version 13_8 (included in the QIIME software package mentioned above), based on the RDP Classifier v2.2, and also by using QIIME-compatible SILVA (8) and BLAST (9). Results indicated that there were no fungi in the beach sand without plastics no alpha and beta diversity statistics were performed. Good’s Coverage was used to estimate the percent of total species represented in the sampling.

## Results

Results of plastics measurements confirmed the transition between contaminated and uncontaminated beach sands (See Table). High throughput DNA sequencing resulted in 80,000-200,000 unique tags, and with >99.9% Good’s Coverage. Analysis indicated that no fungi were present in the beach sand without plastics, thus no alpha or beta diversity analyses could be performed on this dataset. In contrast, the plastic-contaminated beach sand had a fungal community comprised of both free-living and ostensibly plant-associated fungi, dominated by only a few taxa (Figure). Of particular interest are *Acremonium, Penicillium, Malassezia* and *Russula*.

## Discussion

We have indications of at least two free-living fungi that show promise for plastics degradation in beach sands, namely *Acremonium* and *Penicillium. Acremonium* is known to degrade petrochemical contaminants in the lab (e.g., 10). The species of *Acremonium* present here, *A tubakii* is a marine fungus that is known to produce high levels of laccase, an enzyme that is known to degrade plastics (e.g., 11). Similarly, our data support earlier indications that *Penicillium* (e.g., 12) could be a good candidate for plastics breakdown in a wide range of habitats. Both fungi will therefore be put forward for future study by our lab as both free-living solutions to plastics contamination and as enzyme sources. *Malassezia* is a yeast-like pathogenic fungus that causes peritonitis, blood infections and an assortment of chronic and recurrent skin diseases (e.g., 13, 14). While this fungus may be utility as an enzyme source, the human health implications most likely preclude its use in field settings and this result has interesting implications for the impacts of marine plastics on human health.

Interestingly two ectomycorrizal fungi *Russula* and *Inocybe*, were found in the Top 10 most abundant fungi in plastics-contaminated beach sand. Ectomycorrhizal fungi are endosymbionts of most flowering plants and conifers and provide these plant hosts with the nitrogen necessary for survival (15). Some ectomycorrhizal fungi appear to be capable of enzymatic breakdown of at least small polymers in soils as a replacement for host plant carbon in times of need (16, 17). These results imply that some ectomycorrhizal fungi might also be attracted to microplastics as a nutrient source. The anatomy and physiology of *Russula* in particular suggests that species in this genus could prove useful in addressing plastics contamination, particularly when used in combination with host plants. These fungi form a contact exploration type of hyphal network that is known to interact closely with nutrient substrates and to produce enzymes that could be useful in plastics degradation (18). When used in combination with host plants they could be used not only for plastics remediation, but also in combined plastics degradation-reforestation efforts.

To summarize, this preliminary survey provides models for at least two free-living fungi (*Acremonium* and *Penicillium*) that could be useful in plastics remediation efforts and at least one ectomycorrhizal fungal genus (*Russula)* that could be used in combination with native plants for phytoremediation strategies. In addition, a severe human pathogen was apparently attracted to the conditions created by plastics in beach sand, a finding that has interesting health implications for beach goers. This survey was only preliminary and we plan a broader survey across plastics no-plastics transitions in beach habitats at this site comparing sand with and without native pines and willows to further our pool of potential fungal remediation agents.

## Acknowledgements

We thank the non-profit SumOfUs for funding for this project.

## Table and Figure

|  |  |
| --- | --- |
| Microplastics Measures in an Oregon Beach Sand Habitat. |  |
| Beach Without Plastics | 0 |
| Beach With Plastics | 12.6µm +/- 5.5* |
| *Detection range 1-5000µm |  |

**Figure :**
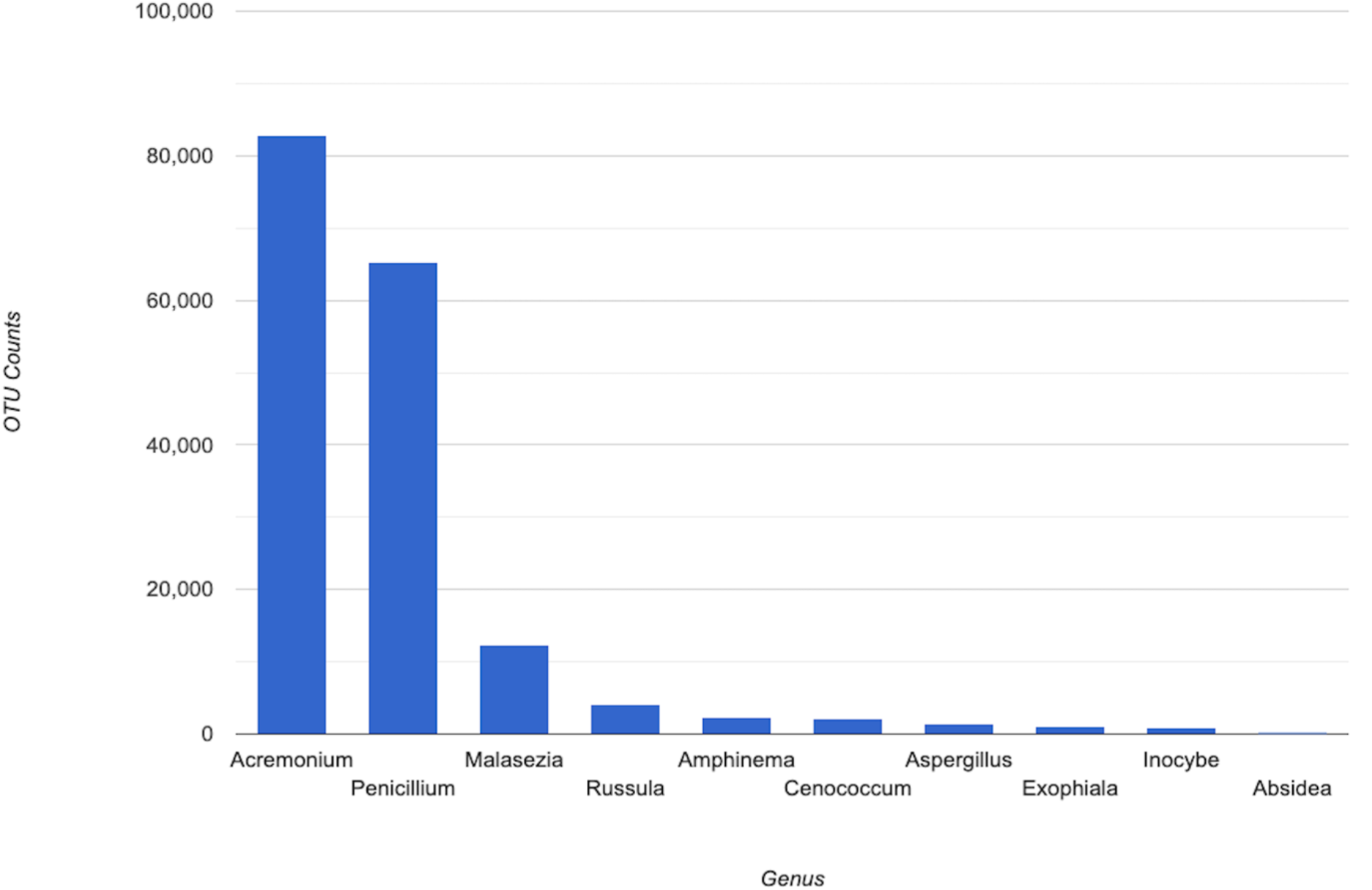
Absolute frequency of top 10 fungal genera in beach sand contaminated by plastic.

